# The impact of radio-tags on Ruby-throated Hummingbirds (*Archilochus colubris*)

**DOI:** 10.1101/008029

**Authors:** Theodore J. Zenzal, Robert H. Diehl, Frank R. Moore

**Affiliations:** Department of Biological Sciences, University of Southern Mississippi, Hattiesburg, Mississippi, USA; U.S. Geological Survey, Northern Rocky Mountain Science Center, Bozeman, Montana, USA

**Keywords:** radio transmitters, radio telemetry, Ruby-throated Hummingbirds, Archilochus colubris, radio-tagging, behavior, flight simulations, migration

## Abstract

Radio telemetry has advanced the field of wildlife biology, especially with the miniaturization of radio-tags. However, the major limitation faced with radio-tagging birds is the size of the animal to which a radio-tag can be attached. We tested how miniature radio-tags affected flight performance and behavior of Ruby-throated Hummingbirds (*Archilochus colubris*), possibly the smallest bird species to be fitted with radio-tags. Using eyelash adhesive, we fitted hatch year individuals (n=20 males and 15 females) with faux radio-tags of three sizes varying in mass and antenna length (220mg-12.7cm, 240mg-12.7cm, and 220mg-6.35cm), then filmed the birds in a field aviary to quantify activity budgets. We also estimated flight range using flight simulation models. When the three radio-tag packages were pooled for analysis, the presence of a radio-tag significantly decreased both flight time (-8%) and modeled flight range (-23%) when compared to control birds. However, a multiple comparison analysis between the different packages revealed that there was a significant difference in flight time when the larger radio-tag package (240mg) was attached and no significant difference in flight time when the lighter radio-tag packages (220mg) were attached. Our results are similar to other studies which analyzed the flight time or flight range of birds wearing radio-tags. Therefore, currently available light weight radio-tags (≤220mg) may be a new option to aid in the study of hummingbird biology. Future study should focus upon the additional drag created by the radio-tag and the effects of the lightest radio-tag packages on free ranging birds. These studies would provide additional information to determine the feasibility on the use of radio-tags to study hummingbird biology.

## Introduction

Radio telemetry has advanced our understanding of wildlife biology in lock step with advances in technology. Using radio telemetry as a way to study birds started in the early 1960s (e.g., Lord et al. 1962; Southern 1964; Graber and Wunderle 1966) and is now widely used to remotely collect movement data of free-ranging birds. Advances in technology have allowed for miniaturization of radio-tags, which has enabled researchers to radio-tag smaller and smaller animals, including hummingbirds (e.g. Hadley and Betts 2009) and arthropods (e.g. Wikelski et al. 2006, 2010). The usual limitation in the use of radio-tags in avian biology is the weight of the radio-tag in relation to total body mass, which is recommended to remain <3-5% (Cochran 1980; Gustafson et al. 1997; Fair et al. 2010). The added weight of a radio-tag may decrease the probability of nesting and increase energetic expenditure (Barron et al. 2010), yet field studies have found that tags up to 5% of the bird’s total body weight do not meaningfully affect survival or movements of small (< 20 g) birds (Naef-Daenzer et al. 2001; Hadley and Betts 2009).

Some research on passerines has shown that radio-tags negatively affect survival (Dougill et al. 2000; Mattsson et al. 2006), while other studies on large, mostly flightless birds suggest negative impacts based on increased energy expenditure (Osborne et al. 1997; Godfrey et al. 2002; Guthery and Lusk 2004). However, most studies investigating direct impacts of radio-tags on survival rates have found negligible effects, if any (Powell et al. 1998; Naef-Daenzer et al. 2001; Hernández et al. 2004; Terhune et al. 2007; Anich et al. 2009; Townsend et al. 2012). The detrimental effects of radio-tags described by Dougill et al. (2000) were due to tag design, while Mattsson et al. (2006) found decreased survival when outfitting nestlings with radio-tags. Long-term survival of radio-tagged birds does not seem to be impeded by tags, as long as tags are well designed and attached after fledging. Additionally, temporary radio-tags, in which the tag eventually falls off, would likely affect survivorship the least (e.g. Raim 1978, Sykes et al. 1990; Naef-Daenzer 1993; Naef-Daenzer et al. 2001; Anich et al. 2009; Hadley and Betts 2009; Smolinksky et al. 2013).

Even if there is no increased likelihood of mortality or reduced reproductive success while wearing a radio-tag, other influences might handicap organisms during particular times of their annual cycle. While Barron et al.’s (2010) meta-analysis on the effects of radio telemetry found no significant effect on flight ability, a radio-tag externally mounted to the back of a bird will necessarily increase body drag (Obrecht et al. 1988; Pennycuick et al. 2012). The main variable in telemetry effect studies is the ratio of the equipment weight to body; arguably the additional drag created by the radio tag has a larger impact on flying animals than the increase in weight. An increase in body drag has been estimated (all things being equal) to reduce long-distance flight ranges, such as during migration (Obrecht et al. 1988; Powell et al. 1998; Pennycuick 2008; Pennycuick et al. 2012). Additionally, the extra weight of a radio-tag may exacerbate energy expenditure of flight, an especially difficult problem for migrating birds that are already carrying increased fat loads. Nonetheless, most investigators make no attempt to determine any impacts of the device before implementing a radio tracking study.

The impact of radio-tags on birds is usually not tested prior to application on free-flying birds, especially for drag. If flight performance is affected by the weight of a radio-tag, then we would expect birds with the heaviest radio-tags to experience the largest effect. Drag, however, varies with transmitter and antenna size, not with mass (Pennycuick et al. 2012). Differing antenna lengths could have a disproportionate impact on the transmitter center of gravity imposing increased energetic costs per unit flight time on individuals outfitted with a longer antenna than individuals outfitted with a shorter antenna. Individuals with the longer antenna will either compensate energetically to the increased flight costs or fly less.

We analyzed the impact of radio-tags on the flight performance and behavior of Ruby-throated Hummingbirds (*Archilochus colubris*), a long-distance, migrant (likely both trans-Gulf and Circum-Gulf) traveling between breeding and wintering destinations (Weidensaul et al. 2013), and the smallest bird to our knowledge to be outfitted with a radio-tag. We used a pairwise study design on individuals in a controlled setting to examine three different radio-tag packages varying in weight and antenna length during fall migration, a time when individuals are accumulating additional mass via fat stores in order to fuel migratory flights. Our two main objectives *a priori* were: 1) quantify the flight time of birds with and without a radio-tag, and 2) estimate the flight range of birds with and without a radio tag from a mechanical model of weight and drag (Pennycuick 2008). A secondary objective *a postori* was to determine if preening behavior differed between treatments.

## Methods

### Study Site and Field Methods

We captured Ruby-throated Hummingbirds using nylon mist nets at the Bon Secour National Wildlife Refuge, Fort Morgan, Alabama (30°10’N, 88°00’W), between sunrise and noon from September 3 – 16, 2010. Netting effort was both active, baiting 10 nets with artificial feeders, and passive. We banded hummingbirds with a USGS aluminum band, aged and sexed them (Pyle 1997), estimated fat (Helms and Drury 1960), measured wing chord and mass, and took a wing photo to determine wing span and wing area for flight range estimates.

### Aviary Routine and Radio-Tag Attachment

We randomly selected a sub-sample of hatch year birds (n=35; mass= 3.80 ± 0.73 g for males (n=20) and 3.76 ± 0.46 g for females (n=15) [these and following results are reported as mean ± SD]). We individually placed birds selected for experimentation into a field aviary (2.43 m X 1.31 m X 1.94 m) with a perch and a feeder without a perch. We used a pairwise study design in which all individuals received, in random order, the control treatment (no radio-tag) and one of three experimental treatments (with faux radio-tag, Figure 1). In each experimental treatment the faux radio-tag varied by antenna length and/or mass. The first experimental group (n=15) received a heavy radio-tag (240mg; total body mass: 6.00% females, 6.32% males) with a long antenna (length: 12.7 cm; diameter: 0.229mm). The second experimental group (n=10) received a light radio-tag (220mg; total body mass: 5.50% females, 5.79% males) with a long antenna (length: 12.7 cm; diameter: 0.152mm). The third experimental group (n=10) received a light radio-tag (220mg; total body mass: 5.50% females, 5.79% males) with a short antenna (length: 6.35 cm; diameter: 0.152mm). Radio-tag design and two faux transmitters were provided courtesy of Sparrow Systems.

**Figure 1.**
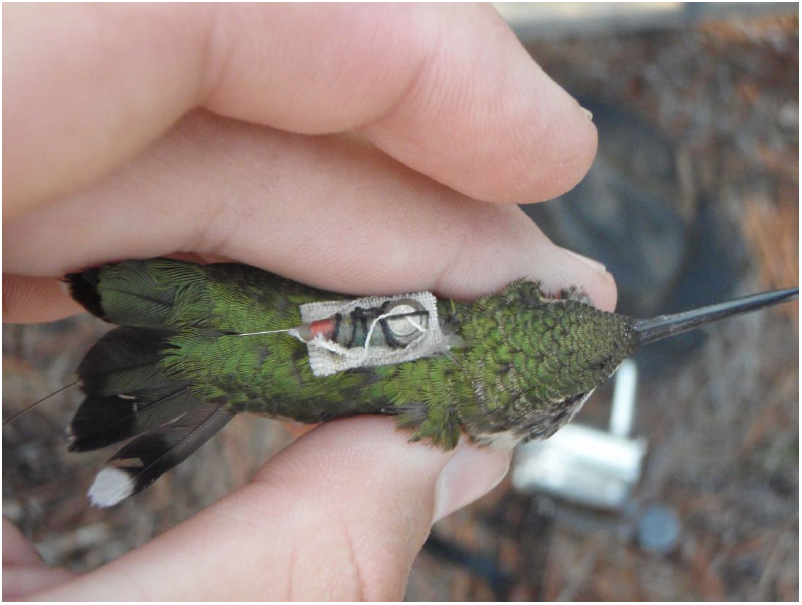
Ruby-throated Hummingbird with a faux radio-tag attached (photo by T.J. Zenzal).

We attached radio-tags using a modified version of Raim’s (1978) method. Radio-tags were first sewn to a piece of cloth the size of the radio-tag, then a second piece of cloth similar in size to the one sewn to the radio-tag was glued to the back of the bird using Revlon^®^ Fantasy Lengths^®^ eyelash adhesive. The cloth sewn to the radio-tag was glued to the cloth on the bird (Figure 1). Cloth, thread, and glue were not included in the radio-tag weight. We removed radio-tags by clipping feathers under the cloth. Treatment order was randomized between individuals to eliminate any effect of order on subsequent analysis. Each individual tested only one radio-tag. After being prepared for the appropriate treatment (with or without radio-tag), individuals were placed in the aviary, allowed to acclimate for 10 minutes before the treatment was recorded, and then videotaped (Panasonic PV-GS65) at 1/4000 frames per second for 7 minutes to score behaviors and time spent in various activities. We then prepped the same individual for the next treatment (attachment or removal of radio-tag), allowed another 10 minutes for acclimation, and then videotaped for another 7 minutes. After a bird completed both treatments, we released it without a radio-tag.

### Flight time

Flight time was quantified from the total 14 minutes (7 minutes for control treatment and 7 minutes for experimental treatment) of video recording. We defined flight as any period an individual was not perched, not distinguishing hovering flight (including feeding) from forward flight. Body condition was determined for two reasons: 1) birds were randomly selected during migration, differing in the amount of fat carried, and 2) this species exhibits reverse sexual-size dimorphism (Weidensaul et al. 2013). We calculated body condition (fat) based on mass and wing length of Ruby-throated Hummingbirds captured on the Bon Secour NWR following Ellegren (1989, 1992) and Owen and Moore (2006). For both sexes, we determined fat free masses by regressing body mass on fat score for individuals in the same wing chord class (1mm increments). The intercept of the regression provided an approximation of fat free mass for a sex-specific wing chord class. After performing regressions on all wing chord classes of individuals included in the study, we executed a second linear regression by regressing the intercepts on each related wing chord class for each sex. The resulting equation from the second regression allowed an estimation of size specific fat free masses for each hummingbird.

Restricted maximum likelihood (REML) models were used to assess the influence of radio-tags on flight time data using JMP statistical software (v.10, SAS Institute 2013). We performed a preliminary analysis on a subset of individuals balanced by experimental order (n = 14 for each group) to determine if the order in which treatments were applied had an effect on flight time. Flight time (square root transformed) was the response variable in a repeated measures mixed model with radio-tag type, experimental order, and presence/absence of radio-tag as fixed factors and individual as a random nested blocking factor. We removed experimental order from subsequent analysis (see Results). We then analyzed flight time data using a repeated measures mixed model with radio-tag type, sex, body condition (described above), and presence/absence of radio-tag as fixed factors and individual as a random nested blocking factor. To determine the impact of each radio-tag type, we reran the analysis for the three radio-tags separately and interpreted p-values using the Holm-Bonferroni adjustment method for multiple comparisons (Holm 1979). Body condition failed to meet the assumptions of normality (Shapiro-Wilk (p=0.02), even after attempting all standard transformations. Therefore, we ran each analysis with and without body condition included as a covariate. To further explore the relationship between body condition and activity budget, we used a Spearman’s rank correlation between flight time and body condition.

### Preening Behavior

We quantified the number of preening occurrences from the video analysis of each bird. Preening is defined as each time an individual’s bill made contact with its feathers. We analyzed preening occurrences using a generalized linear mixed-effects model fit by maximum likelihood with a quasi-Poisson distribution (O’Hara and Kotze 2010) in the R statistical language (R Core Team 2013), package “MASS” (Venables and Ripley 2002). The fixed factors of this model included radio-tag type, experimental order, body condition, and presence/absence of radio-tag, with individual as a random nested blocking factor. We excluded sex in this model since we did not believe there is any biological significance of sex on preening. We did however include experimental order because observations suggested that birds receiving the experimental treatment first may preen more after the back feathers were clipped to remove the transmitter. To further explore interactions we performed two additional tests in the R statistical language: First, a Nemenyi post-hoc test (Hollander and Wolfe 1999) from package “coin” (Hothorn et al. 2006, 2008) was performed on significant effects of the model. Second, a Spearman’s rank correlation was used to determine a relationship between preening occurrences and flight time.

### Flight Range Estimates

We used Program Flight 1.24 to estimated flight range of birds with and without radio-tags. Simulations were based on wing area, fat free mass, and body condition (Pennycuick 2008), for each bird (n=31) with and without a radio-tag. We obtained fat free masses and body conditions of Ruby-throated Hummingbirds using methods described above. However, three of the individuals fell below the average fat free mass and according to the conditions of the model were not able to migrate. Therefore, two females and one male were eliminated from analysis due to lean body condition; additionally, a third female was removed because she had no associated wing photograph. We quantified wing span and wing area as described in Pennycuick (2008), although we photographed rather than hand traced wings. We modified Pennycuick’s (2008) wing area quantification by using ImageJ (Abramoff et al. 2014) to determine the exact wing area (partial wing area plus rootbox) from a digital tracing of an individual’s semi-span instead of using a grid to quantify area as performed with hand tracings. We assumed trans-Gulf flight in still-air conditions and an altitude of 500 m (air density of 1.17 kg/m^3^; Kerlinger and Moore 1989; Woodrey and Moore 1997). When an individual was simulated with a radio-tag, a payload mass was determined for the appropriate radio-tag size (220mg or 240mg) with a drag factor of 1.5 (Pennycuick et al. 2012). We determined differences between simulated flight ranges (square root transformed) for individuals with and without a radio-tag using a REML repeated measures mixed model. Radio-tag type, sex, and presence/absence of radio-tag were set as fixed factors and individual as a random nested blocking factor. This statistical analysis was performed using JMP statistical software (v.10, SAS Institute 2013).

## Results

### Flight time

We found mixed evidence of experimental order impacting flight time of hummingbirds, due to a significant interaction between experimental order and presence/absence of a radio-tag (F_1,55_=21.12, p=0.0001). Individuals decreased flight time during the second treatment regardless of treatment order. There was, however, much individual variation which clouds the interpretation of the results but illustrates that attachment of a radio-tag will not elicit the same response from every individual. Individuals receiving the control treatment first had an 80.00±167.59s decrease when the radio-tag was attached, while individuals with a radio-tag attached first had a 5.95±148.26s decrease during the control treatment. A bird undergoing the control treatment second had feathers clipped which possibly explains why there was decreased flight time during the control treatment. The decrease in the subsequent treatment is likely a result of preening and possibly acclimation to captivity (see below). However, the main effect of experimental order did not affect flight time (F_1,55_=0.32, p=0.58). Based on the main effect test, large amount of inter-individual variation, and the *a priori* effort to randomize treatment order, we concluded that experimental order did not meaningfully impact activity budgets and excluded it as a factor from the subsequent analysis.

Flight time was about 8% less with a radio-tag attached (F_1,69_=7.36, p=0.01 without body condition as a factor; F_1,69_=6.00, p=0.02 with body condition as a factor). Flight time without a radio-tag (182.94 ± 121.72s) was greater than when a radio-tag was attached (149.6±104.39s, averaged across all models). However, size of the radio-tag did not have a significant effect on flight time in either model (F_2,69_=0.98, p=0.39 without body condition as a factor; F_2,69_=1.83, p=0.18 with body condition as a factor). Further analysis using multiple comparison testing between radio-tag types using the Holm-Bonferroni adjustment (Holm 1979) revealed that the only radio-tag to have a significant decrease (∼11%) in flight time between the treatment and the control was the heavy tag (F_1,29_=15.06, p=0.002 without body condition as a factor; F_1,29_=13.27, p=0.004 with body condition as a factor; adjusted α=0.017), while flight time in both of the light tag treatments did not differ from controls (long antenna tag: ∼8% decrease, F_1,19_=1.76, p=0.22 without body condition as a factor; F_1,19_=2.32, p=0.18 with body condition as a factor; adjusted α=0.025; short antenna tag: ∼6.5% increase, F_1,19_=0.05, p=0.83 without body condition as a factor; F_1,19_=0.06, p=0.82 with body condition as a factor). Mean flight time decreased predictably from the light weight short antenna tag having the most flight time (210.70±127.14s), followed by the light weight long antenna tag (148.40±74.93s), while the heavy weight long antenna tag had the least flight time (137.00±107.92s)(Figure 2). Although wing morphology of Ruby-throated Hummingbirds is sex-dependent (Stiles et al. 2005), neither model showed an effect of sex on flight activity (F_1,69_=2.31, p=0.14 without body condition as a factor; F_1,69_=1.24, p=0.28 with body condition as a factor) nor an interaction between sex and the presence of a radio-tag (F_1,69_=0.21, p=0.65 without body condition as a factor; F_1,69_=1.49, p=0.24 with body condition as a factor). Finally, body condition did not impact flight time (F_1,69_=1.35, p=0.26), and a Spearman’s rank correlation showed no relationship between flight time and body condition (with radio-tag Spearman’s rho= −0.01, p=0.94, n=35; without radio-tag: Spearman’s rho= −0.18, p=0.30, n=35).

**Figure 2.**
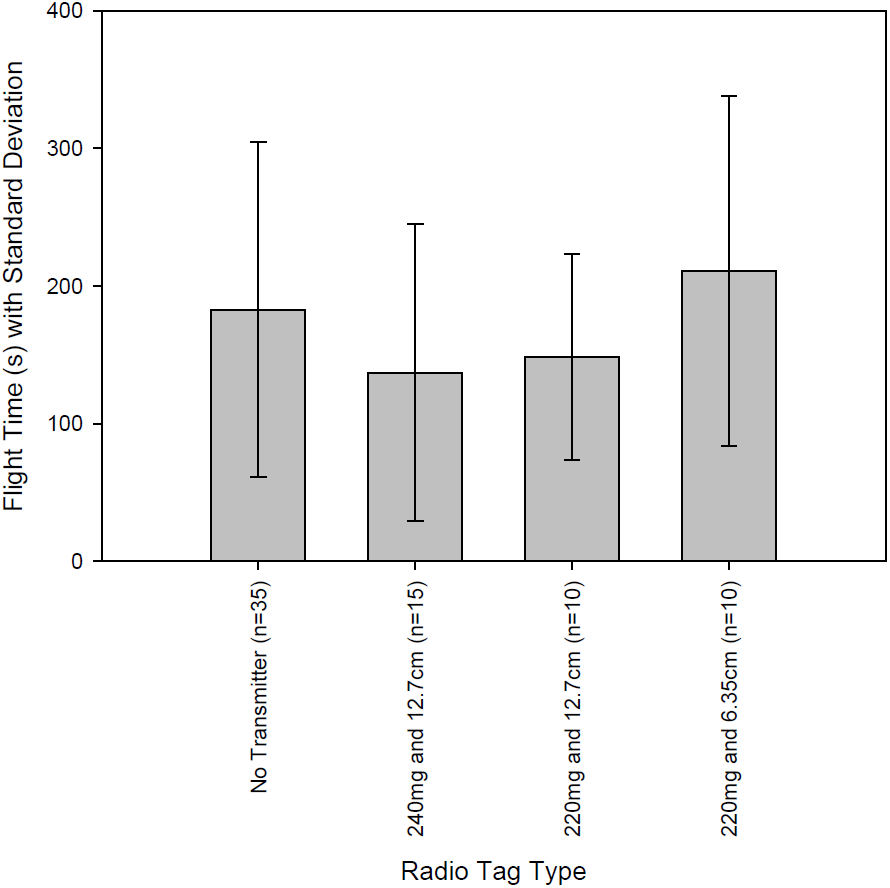
Mean flight time (s) of Ruby-throated Hummingbirds per 7 min treatment with (240mg: n=15; 220mg: n=10) and without (n=35) radio-tags attached. Radio-tags are separated by mass (mg) and length of antenna (cm). Standard deviation is shown as vertical bars.

### Preening Behavior

As expected, initial analysis revealed that preening occurrences had a significant negative correlation with flight time (with radio-tag: Spearman’s rho= −0.39, p=0.02, n=35; without radio-tag: Spearman’s rho=−0.38, p=0.02, n=35), which is not surprising given preening and flying are mutually exclusive behaviors. Preening occurrences did not differ between the presence/absence of a radio-tag (t=0.18, df=23, p=0.86, n=35). However, experimental order did have a significant effect on the number of preening occurrences (t=2.43, df=23, p=0.02, n=35). Birds receiving the control treatment first (n=16) had a mean of 25.31±42.53 preening occurrences which increased to 37.63±55.15 preening occurrences when the radio-tag was attached. Birds that first received the experimental treatment (n=19) increased preening occurrences from 27.84±35.06 to 30.26±40.71 after the radio-tag was removed. However, given the close means and significant effect of preening on experimental order, a Nemenyi post-hoc test (Hollander and Wolfe 1999) was performed on the number of preening occurrences by order which yielded a non-significant effect (Z=0.31, p=0.76, n=35). Although the number of preening occurrences does not differ significantly while an individual has a radio-tag attached, the number of preening occurrences are greater when a radio-tag is attached (32.31±35.38) compared to when no radio-tag is attached (28.00±41.01) averaged across treatments. There is much individual variation between subjects regardless of treatment. When a radio-tag was attached preening occurrences ranged from 0 to 184 instances over 7 minutes, individuals without a radio-tag attached showed a similar range from 0 to 157 instances over the same time frame. This analysis further explains the significant interaction between presence/absence of a radio-tag and experimental order for flight time.

### Flight Range Estimates

Flight modeling provided an estimate of how a radio-tag affects flight range. The presence of a radio-tag significantly affected simulated flight ranges (F_1,62_=135.26, p<0.0001), reducing an individual’s flight range by ∼340 km on average (without radio-tag: 1512.26±1188.65 km; with radio-tag: 1172.00±916.86 km). There was no effect of sex on flight range (F_1,62_=0.48, p=0.49), nor was there an effect of radio-tag mass (F_2,62_=1.08, p=0.36).

### Discussion

Hummingbirds are challenging to study because their size and speed of movement makes detection of birds difficult by visual means. The ability to continually track and record the behavior of radio-tagged hummingbirds would measurably enhance our understanding of migratory movement, dispersal, resource use, home range activity, and habitat use. For example, the first published application of a radio-tag on a hummingbird determined the movement patterns of Green Hermits (*Phaethornis guy*) in Costa Rica (Hadley and Betts 2009). To our knowledge, the Green Hermit is the smallest bird (5.8±0.09 g; Hadley and Betts 2009) that has been radio-tagged prior to our study of Ruby-throated Hummingbirds. The miniaturization of transmitters has allowed others to track flying arthropods, much smaller than most hummingbirds, which provided insight to questions that would be difficult to answer using other means (e.g., Wikelski et al. 2006, 2010; Pasquet et al. 2008; Hagen et al. 2011).

Activity budgets of Ruby-throated Hummingbirds are influenced by the presence of a radio-tag, although only the heaviest radio-tag showed a significant decrease (∼11%) in flight time from the control treatment. The light radio-tags had less influence in flight time with the long antenna tag decreasing flight time by ∼8%, and the short antenna tag increasing flight time by ∼6.5%. These radio-tags at 220mg, just over 5% total body mass of a Ruby-throated Hummingbird, did not seem to pose a significant handicap on activity, similar to other studies using radio-tags at a comparable percent body weight (Naef-Daenzer et al. 2001; Hadley and Betts 2009). However, it is difficult to extrapolate the small differences found in flight activity that might be an artifact of a seven minute experimental period in an aviary to actual migratory flight.

Free ranging animals may behave differently when in captivity (see Clubb and Mason 2003). The size of the aviary or simply being placed in an aviary may have limited the activity of the hummingbirds once they determined there was no way out. Although time of day was not included as a factor in analysis, most birds were tested in the late morning or early afternoon when they are typically inactive (Zenzal, personal observation) or migrating (Hall and Bell 1981; Willimont et al. 1988). The length of time allotted for birds to acclimate to the radio-tag may have been too short, affecting the outcome; most birds receiving any sort of marker (e.g. band, radio-tag) spend an unpredictable amount of time reacting to the tag (preening or attempting to remove the marker) before resuming normal behaviors.

Preening increased in individuals that received the experimental treatment compared to the control treatment. Increases in comfort behavior (as described by Delius 1988; i.e. preening, wing flapping, head shaking) would necessarily increase the amount of time spent perching, while decreasing time spent in flight. While preening explains some of the variation found during flight activity, particularly between the different experimental orders which may be due to attaching radio-tags directly to feathers or clipping back feathers to remove the radio tag, caution is recommended when making interpretations from this analysis as handling birds seemed to increase the likelihood of birds preening. Although we found no significant effect of a radio-tag on preening, other studies have shown that preening did increase with the attachment of a transmitter (Hooge 1991; Pietz et al. 1993; Sykes et al. 1990).

The apparent effect of the radio-tag on activity might be influenced by attachment method, since the radio-tag was glued directly to back feathers instead of skin for easy removal after the experiment was complete. The most common adhesive attachment method requires feathers to be clipped in order to create a strong bond between the radio-tag backing and the skin of the bird or feather shaft (e.g., Raim 1978; Sykes et al. 1990; Naef-Daenzer 1993; Naef-Daenzer et al. 2001; Anich et al. 2009; Hadley and Betts 2009; Smolinksky et al. 2013). The effects of this attachment method are negligible on small birds compared to other attachment methods tested (Sykes et al. 1990). Furthermore, field studies showed no decrease in survivorship when this attachment method was used compared to non-radio-tagged birds (Naef-Daenzer et al. 2001; Anich et al. 2009).

The percent body mass and size of radio-tags, but not the antenna length, appeared to affect the activity budget; however much variation existed between and within treatments. Our findings are consistent with the influence of drag of the device (Barron et al. 2010; Pennycuick et al. 2012) rather than the weight of the radio-tag viz. energetic expenditure. The light radio-tag with the short antenna had the highest amount of flight time, likely due to decreased drag of the antenna. However, large individual variation across all the variables explored make it difficult to suggest any hard-and-fast rules for radio-tagging hummingbirds, besides selecting a radio-tag that has the smallest weight and drag available. A valuable follow-up study to the one described would determine the amount of drag different radio-tag designs have on free-flying hummingbirds, similar to Pennycuick et al.’s (2012) study of external device drag on Rose-coloured Starlings (*Pastor roseus*), with the use of a wind tunnel.

Predicted flight range was affected by the presence of a radio-tag but did not vary with size of the radio-tag or sex of the individual. Individuals able to fly farther by virtue of larger fat loads experienced larger decreases in distance when radio-tagged compared to individuals with shorter flight ranges. Although flight simulations showed a decrease in flight ranges, most studies on survival and return rates of radio-tagged long-distance migrants fail to show an effect of radio-tags on survival (Powell et al. 1998; Cardinal 2005; Anich et al. 2009; Townsend et al. 2012; however see Samuel and Fuller 1996). In two of these studies, a subsample of birds tagged were recaptured a year later (in one case 2 years later, Powell et al. 1998) with radios still attached (Powell et al. 1998; Townsend et al. 2012).

Hummingbird behavior is potentially affected by the presence of a radio-tag, so caution should be exercised when selecting individuals to tag, which will depend on season, sex, and condition. For example, a radio-tag is likely to impede nest construction in female hummingbirds (see Weidensaul et al. 2013). That said, observations of free flying Ruby-throated Hummingbirds during stopover revealed that individuals with radio-tags behave similarly to marked individuals without radio-tags in stopover duration, foraging, competitive interactions, and seasonally appropriate departure directions (Zenzal, personal observation). One of these free flying radio-tagged birds was detected, wearing its tag, at an artificial feeder in Corpus Christi, Texas (∼950 km from Fort Morgan, Alabama) two weeks after being tagged (USGS Bird Banding Laboratory, personal communication).

## Acknowledgments

Research funding was provided by the Department of Biological Sciences at the University of Southern Mississippi (USM) and a grant from the National Science Foundation (NSF) to FRM (IOS 844703). TJZ was also supported by a fellowship from the NSF GK-12 program “Molecules to Muscles”, Award # 0947944 through USM. We would especially like to thank the Bon Secour National Wildlife Refuge and Fort Morgan Historic Site for allowing us to work on their properties. We would also like to thank the 2010 Fort Morgan Field crew for help with data collection, C.J. Pennycuick for insightful conversations during the formulation of this manuscript, C. Qualls and J. Schaefer for statistical advice, and J. Cochran for conversations about designing radio-tags for hummingbirds and supplying faux radio-tags. We would also like to thank the Migratory Bird Research Group at USM for their support. Finally, we would like to thank Z. Németh and anonymous reviewers for helpful comments on this manuscript. This study was approved by the USM Institutional Animal Care and Use Committee as well as the USGS Bird Banding Laboratory. Any use of trade, firm, or product names is for descriptive purposes only and does not imply endorsement by the U.S. Government.

